# Effects of the maternal factor Zelda on zygotic enhancer activity in the *Drosophila* embryo

**DOI:** 10.1101/385070

**Authors:** Xiao-Yong Li, Michael B. Eisen

## Abstract

The maternal factor Zelda is broadly bound to zygotic enhancers during early fly embryogenesis, and has been shown to be important for the expression of a large number of genes. However, its function remains poorly understood. Here, we carried out detailed analysis of the functional role of Zelda on the activities of a group of enhancers that drive patterned gene expression along the anterior -posterior axis. We found that among these enhancers, only one lost its activity entirely when all its Zelda bind sites were mutated. For all others, mutations of all of their Zelda binding sites only had limited effect, which varied temporally and spatially. These results suggest that Zld may exert a quantitative effect on a broad range of enhancers, which presumably is critical to generate highly diverse spatial and temporal expression patterns for different genes in the developmental gene network in fly embryo. Lastly, we found that the observed effect of Zelda site mutations was much stronger when a mutant enhancer was tested using a BAC based reporter construct than a simple reporter construct, suggesting that the effect of Zld is dependent on chromatin environment.

## Introduction

Enhancers that drive patterned gene expression during animal development are usually bound by multiple transcription factors (Lelli et al., 2012; Spitz and Furlong, 2012). Most of these factors directly provide temporal and spatial cues and are responsible for the precise expression patterns of their target genes. However, a group of factors, termed the pioneer factors have been shown to play a predominant role in determining transcription factor access to enhancers in closed chromatin and the overall enhancer activity (Zaret and Carroll, 2011).

Owing to their importance in enhancer function, it will be important to gain better understanding about the functions of these factors. In this regard, previous studies have indicated that pioneer factors can play diverse functional roles through out development. They have been suggested to be important in determining the temporal dynamics of target gene expression (Gaudet and Mango, 2002), development enhancer prepatterning in cell lineage specification (Boller et al., 2016; Gualdi et al., 1996; Heinz et al., 2010; Mayran et al., 2018; van Oevelen et al., 2015; Xu et al., 2011), and in maintaining functional competence of developmental enhancers by preventing the formation of repressive chromatin(Xu et al., 2009), among others.

Zelda(Zld) is a maternal factor which is essential for zygotic genome activation in flies(Liang et al., 2008). It recognizes the CAGGTAG motif, which was found to be broadly associated with zygotic gene promoters and enhancers in early fly embryos (Bosch, 2006; Li et al., 2008; Nègre et al., 2011; Satija and Bradley, 2012; Yáñez-Cuna et al., 2012), and not surprisingly, has been found to bind broadly in early embryos, (Harrison et al., 2011; Nien et al., 2011). Since Zld has been found to bind to zygotic enhancers as early as mitotic cycle 8 before most of the zygotic enhancers become active, it has been suggested to function as a pioneer factor (Harrison et al., 2011). In support of this, Zld has been shown to be broadly required for chromatin accessibility at zygotic enhancers (Li et al., 2014)Foo:2014bh, (Schulz et al., 2015).

Several studies have investigated the effect of Zld on zygotic gene expression and enhancer activity(Combs and Eisen, 2017; Foo et al., 2014; Harrison et al., 2011; Liang et al., 2008; Nien et al., 2011; Sun et al., 2015; Xu et al., 2014). However, a detailed understanding of Zld function is still limited. Here we extend previous studies by carrying out detailed analysis on the effect of Zld binding site mutations on a group of anterior-posterior (A/P) enhancers.

## Results

We used transgenic reporter assay to investigate the effect of Zld binding site mutations on a selected group of enhancers. Common enhancer reporter constructs usually contain enhancer sequences placed immediately upstream of a core promoter, which drives the expression of a reporter gene. To assay the enhancers in a more natural setting, we cloned each enhancer 860 bp upstream of the *eve* promoter that drives the *lacZ* reporter gene (Fig. 1A)

**Fig. 1.**
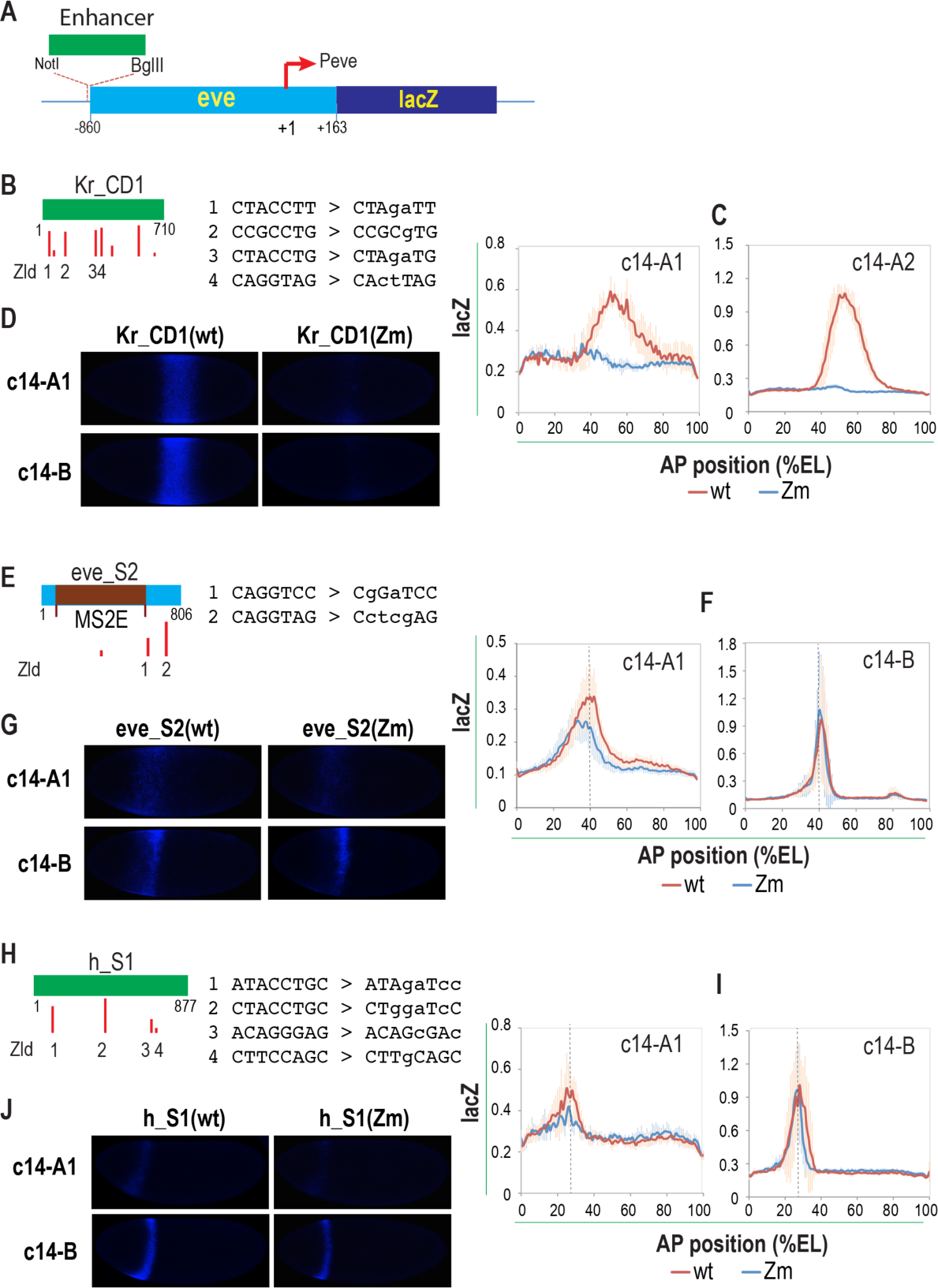
Effect of Zld binding site mutation on the activities of anterior enhancers. A. Schematic of the enhancer reporter constructs created with the pLEP-BFYZ vector. The enhancer sequences were cloned upstream of the *eve* gene sequence from −860 - +163, which includes the *eve* promoter that drives the expression of the *Lac Z* reporter. B, E, H) schematic for each enhancer Is shown. The heights of the red bars represent the weight matrix scores for the Zld binding sites. The sequences of wild type and mutated Zld sites are also shown. The *eve* stripe 2 enhancer sequence used in this study is longer than the previously defined minimal enhancer (MS2E) as indicated. The strong Zld site on the right in the Kr_CD1 enhancer was not mutated since it is located outside the Zld bound region based on ChIP-seq (Harrison et al., 2011). C,F,I) The reporter activities for the wild type (wt) and Zld binding site mutant (Zm) enhancers along the anterior – posterior axis at the indicated stages are shown. The *lacZ* reporter activity in embryos of the transgenic flies was detected by FISH using a *lacZ* antisense probe, and the *lacZ* signal across the A/P axis was quantified and normalized with nuclei density, as described in Material and Methods. D,G,J) The representative embryo images showing the reporter expression in embryos from the transgenic flies for the wild type and mutant enhancer constructs.

We tested five enhancers that are known to drive patterned gene expression along the *A/P* axis. Among them, three drive expression in the anterior part of the embryos and are known to be dependent on Bcd for activation, which include the stripe 1 enhancer, h_S1, of the pair-rule gene *hairy* (Riddihough and Ish-Horowicz, 1991), the stripe 2 enhancer, eve_S2, of the pair-rule gene *even-skipped* (Small et al., 1992), and the CD1 enhancer, Kr_CD1, of the gap gene *Krüpple* (Hoch et al., 1990). The remaining two enhancers drive expression in the posterior part of the embryo including the upstream enhancer tll_K10 of the terminal gene *tailless* (Rudolph et al., 1997), which is activated by torso, and the stripe enhancer, hb_stripe, of the gap gene *hunchback* (Margolis et al., 1995; Perry et al., 2012), which is activated Caudal (Cad) respectively. All these enhancers are strongly bound by ZLD based on previous ChIP-seq analysis (Harrison et al., 2011).

We performed fluorescence *in situ* hybridization (FISH) on fixed embryos collected from transgenic flies created with the reporter constructs using an anti-sense probe for the *lacZ* reporter (Fig. 1A). The embryos were imaged by confocal laser scanning microscopy. The reporter signal along the *A/P* axis was calculated from the images and normalized by the nuclei signal from Sytox Green nucleic acid stain, as described in Materials and Methods. All analyses were carried out in cycle 14 (c14) embryos. We carefully staged the embryos so that the dynamics in the changes of the reporter activity in the mutant and wild type embryos can be compared. We divided the embryos into four age groups, c14-(A–D), based on the extent of cell membrane invagination, and further divided c14-A into c14-A0, A1 and A2 based on nuclear shape as detailed in Materials and Methods.

### Effect of mutating Zld binding sites on anterior Bcd-dependent enhancers

Among the three anterior enhancers, only Kr_CD1 showed near complete loss in activity after all its Zld binding sites were mutated (Fig. 1B,C,D, Fig. S1). This finding is in consistent with the near complete loss of its activity in *zld-* mutant embryo reported in previous study (Xu et al., 2014).

The other two anterior enhancers, h_S1 and eve_S2 behaved very similarly to each other both in terms of their expression pattern dynamics and how Zld binding site mutations affect their activities (Fig. 1E-J, Fig. S2,3). Both enhancers drove the reporter gene expression with a stripe pattern, as expected. The stripe in each case was broad at the start of c14, and became much narrower as the embryos enter mid c14 (Fig. 1F,G,I,J). For both enhancers, when all their Zld binding sites were mutated, there was significant decrease in the reporter activity in early c14 (A0, A1). In older embryos, however, the effect of Zld site mutations became much weaker even though the stripes remained consistently narrower for both mutant enhancers compared to the wild type enhancers. Interestingly, in early c14-A0 and A1, the effect of Zld site mutations was stronger in the posterior part of the stripe than in the anterior half of the stripe, which is more apparent for the eve_S2 enhancer. This differing positional effect of Zld may be explained by the lower Bcd concentration in the posterior part of the expression pattern, which makes the enhancer more dependent on Zld. In support of this interpretation, a previous study has shown that enhancers that are active further away from the anterior of the embryo tend to be more dependent on Zld due to lower Bcd concentration (Xu et al., 2014). Take together, after mutating all their Zld binding sites, the h_S1 and eve_S2 enhancers only experience significant decrease in activity in early c14, and very small change in older embryos. These results differ from effects observed for endogenous *h* and *eve* expression in *zld-* mutant embryos, in which the *h* stripe 1 and *eve* stripe 2 expression were completely abolished (Nien et al., 2011), which is likely due to secondary effects resulting from the lose of Zld in the embryo (see below).

### Effect of mutating Zld binding sites on the posterior enhancers

The tll_K10 enhancer drives strong expression in the posterior cap of the embryo as well driving a modest level of expression in the anterior pole. Zld binding site mutations had no apparent effect of anterior activity. In the posterior, the tll_K10 mutant enhancer with all its Zld binding sites mutated displayed significantly lower activity in early c14 (Fig. 2A-C, Fig. S4). However, the effect of Zld site mutations became very small as the embryo develops further, similar to h_S1 and eve_S2 enhancers. These observed effects of Zld binding site mutation on tll_K10 enhancer activity, again, is not what would be expected based on the behavior of the endogenous *tll* gene in the *zld-* mutant embryo, in which it lost all the expression on the dorsal side of the posterior cap, and the expression expanded toward the anterior in the ventral side of the cap (Nien et al., 2011), presumably due to secondary effect.

**Fig. 2.**
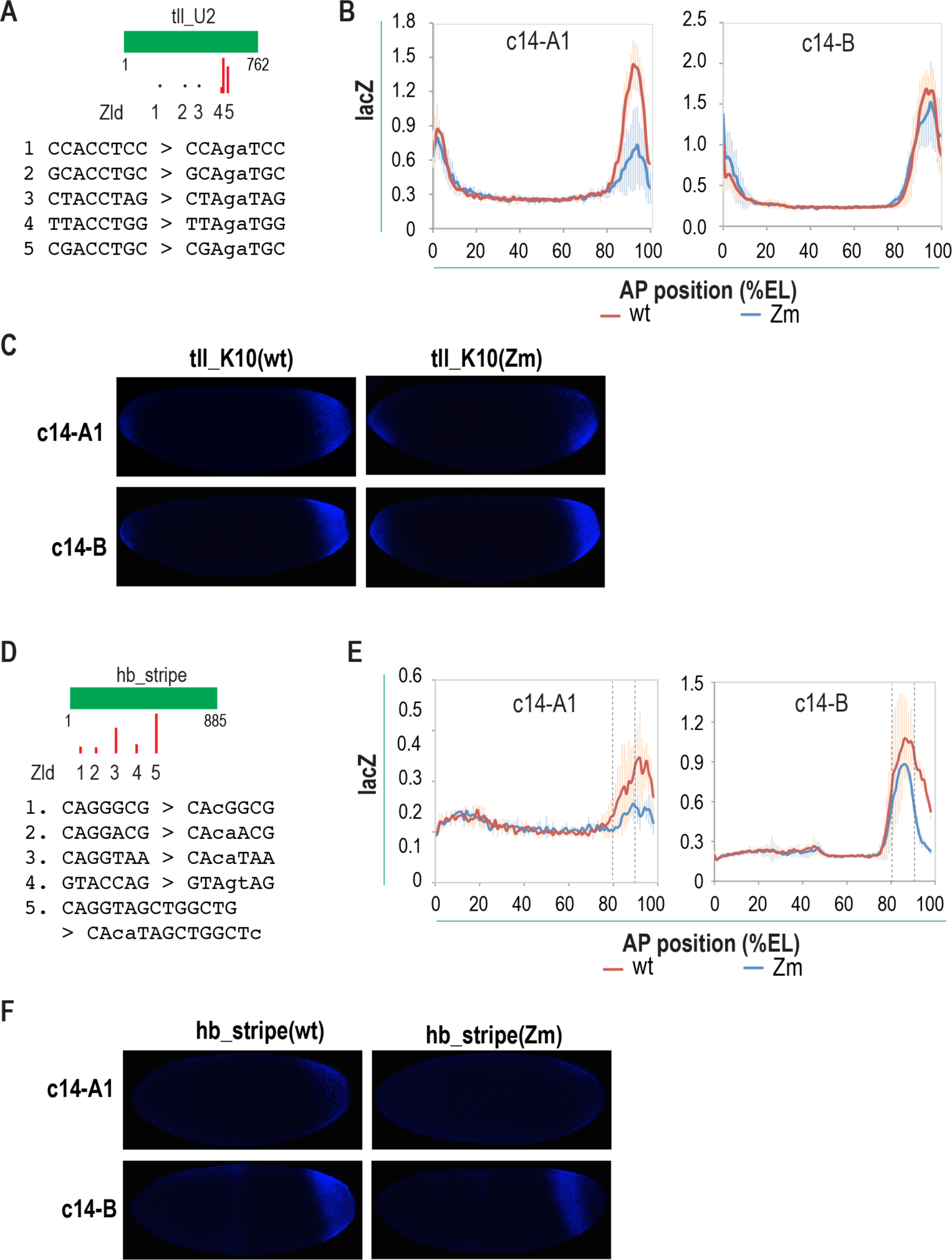
Effect of Zld site mutations on the activities of posterior enhancers. The enhancers were cloned into the pLEP-PBFYZ vector and transgenic lines were created for the resulting constructs, as shown in Fig.1A. The *lacZ* reporter activity was detected, analyzed as described in Fig.1B. A,C) The Zld site mutations for tll_K10 and hb_PS enhancers are shown. B,D) The activities of the wild type and mutant enhancers in embryos at the stages as indicated are shown. C,E) Representative images showing the reporter activities in embryos from transgenic flies containing the wild type and Zld site mutant enhancer constructs.

The *hb* stripe enhancer primarily drives expression in the posterior part of the embryo with a relatively weak middle stripe expression. Here we only examine the effect of Zld binding site mutation on the expression in the posterior. Zld binding site mutations on this enhancer had a complex effect, showing both spatial and temporal variation (Fig. 2D-F, Fig. S5). At the start of cycle 14 (A0 and A1), the activity of the wild type enhancer was detected in a domain between 80-100% egg-length, which is expected based on previous analysis of this enhancer using *hb* BAC plasmid. However, at this time, the mutant enhancer with all its Zld sites mutated showed very little activity, reflecting strong effect of the Zld site mutations. Near the end of c14A, the wild type enhancer displayed strong activity in the same domain between 80%-100% egg length. In comparison, the mutant enhancer displayed near wild type activity between 80–90% egg length, but much lower activity between 90-100% egg length. In late c14, the reporter activity of the wild type enhancer retracted from the posterior, and the mutant enhancer displayed activity that is overall lower than the wild type enhancer.

Much of this varied effect of Zld binding site mutations on the stripe enhancer activity may be explained by the dynamics in the distribution of the Cad protein. Based on immunochemistry (Macdonald and Struhl, 2004), Cad forms a broad posterior-anterior gradient in the pre-cellular blastoderm embryo; starting from the onset of c14, the Cad gradient begins to retract both from the anterior and the posterior but the Cad concentration in a stripe region at about 90% egg length remained almost constant. The relatively stable high concentration in the 80-90% egg length region throughout c14 can explain why the enhancer showed at most modest dependence on Zld in this region, while the fast decline of Cad concentration offers the explanation why the enhancer is more affected by the Zld site mutation in the posterior. The dynamic in Cad distribution, however, does not explain the overall slow appearance of the mutant enhancer activity at the start of c14, which is probably due to the requirement of Zld for faster restart of transactivation following mitosis in general. Taken together, the dependence of the Cad activated stripe enhancer on Zld can mostly be explained similarly by the gradient boosting model used to explain the differing requirement of Zld for Bcd dependent enhancers that are active at different positions along the Bcd gradient.

### Stronger effect of Zld observed using a BAC reporter construct

The relatively modest effects of the Zld binding site mutations on the enhancer activities described above were surprising. Since Zld has been suggested to potentiate transription factor binding by increasing chromatin accessibility at enhancers (Foo et al., 2014; Li et al., 2014; Schulz et al., 2015; Sun et al., 2015), it is possible that the favorable chromatin environment around the enhancer-reporter sequences in the transgenic flies may have alleviated the dependence of the enhancers on Zld. To investigate this possibility, we decided to test the effect of Zld binding site mutations on the *eve* stripe 2 activity using an *eve* BAC reporter construct.

We started with a eve BAC plasmid that contained the full *eve* gene including all its regulatory sequences as well as the two insulators that flank the *eve* locus, which are important to prevent the native chromatin state in the eve locus to be influenced by the flanking chromatin environment. We then modified the BAC plasmid by replacing the *eve* coding sequence in the BAC with the *eGFP* reporter gene, and by mutating all the Zld sites around the *eve* stripe 2 enhancer (Fig. 3A). Transgenic flies were created using this mutant enhancer containing construct as well as the wild type counterpart, and the reporter activities in the embryos from these flies were tested by FISH using an antisense probe for the *eGFP* reporter.

**Fig. 3.**
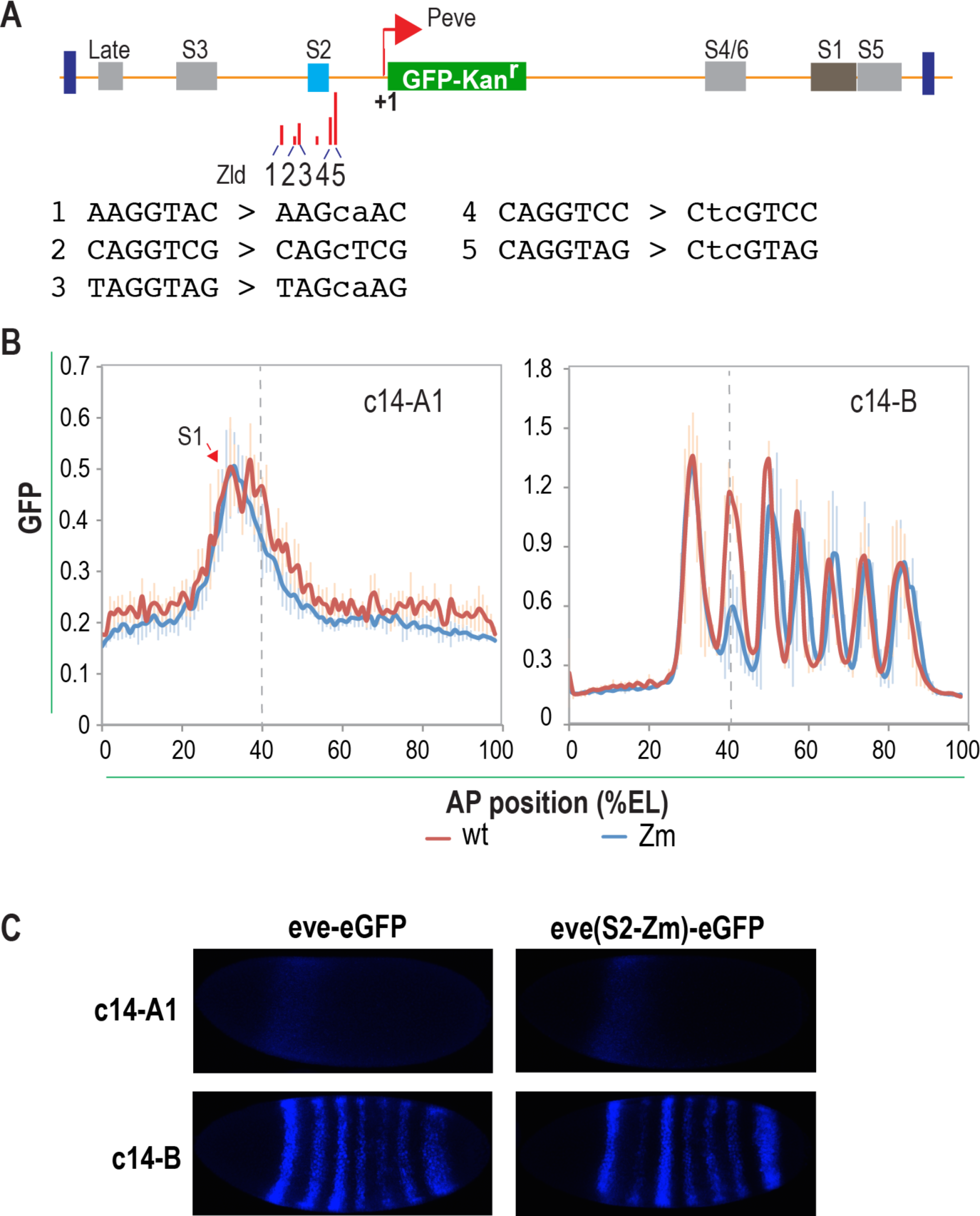
Effect of Zld binding site mutations on *eve* stripe 2 enhancer activity detected using a eve BAC reporter construct. A). Schematic for the eve BAC construct, in which the *eve* CDS is replaced by the *eGFP* reporter gene. The Zld site mutations introduced to the stripe 2 enhancer region in the mutant construct are also shown. B). The *eGFP* reporter activity profiles along the anterior – posterior axis in the embryos from transgenic flies containing the either the wild type construct, or the construct containing Zld binding site mutations in the stripe 2 enhancer region are shown. The reporter activities were detected by FISH and analyzed as described in Fig. 1B except that the eGFP anti-sense probe was used. C) Representative images showing the effect of Zld site mutations on the *eve* stripe 2 enhancer activity.

As shown in Fig.3B and C, at start of c14, the wild reporter construct drove the reporter expression in a broad stripe in the anterior part of the embryo. Since both the stripe 1 and 2 enhancers contribute to this initial expression, it is impossible to assess the magnitude of the effect of Zld site mutations on the stripe 2 activity, even though there was a clear decrease in the activity in the posterior part of this early stripe, which may be attributed to the stripe 2 Zld binding site mutations. However, in older embryos when the two stripes became distinct, a strong effect of the Zld site mutations on the stripe 2 activity became apparent. This observed effect is much stronger than what we saw with the simpler reporter assay described earlier. This suggests that Zld can display different effect under different assay conditions. Nevertheless, since there was still some *eve* stripe 2 expression remaining, this result still does not explain the complete loss of *eve* stripe 2 expression in *zld-* embryos. In addition, in *zld-* embryos genes also often displayed changed expression patterns (Nien et al., 2011; Xu et al., 2014), many of the effects observed in the mutant embryos are likely due to secondary effect.

## Discussion

Several previous studies have investigated the effect of Zld on zygotic gene expression either using genomic methods or *in situ* hybridization for individual genes in *zld-* mutant embryos, which revealed broad dependence of zygotic genes on Zld (Combs and Eisen, 2017; Liang et al., 2008; Nien et al., 2011; Sun et al., 2015; Xu et al., 2014). Additionally, several studies have also investigated the role of Zld on enhancer activity by analyzing the activities of enhancers in *zld-* mutant embryos or the effect of Zld binding site mutations on selected enhancers {Bosch:2006fx, (Foo et al., 2014; Xu et al., 2014). These studies have been useful, but a detailed understanding on the role of Zld in enhancer function has remained limited. Our current study was aimed at extending these previous studies to provide further insight on Zld function.

From this study, we have found that even though the enhancers we chose for our study are among the strongest bound by Zld based on ChIP-seq (Harrison et al., 2011), the binding site mutations for all but one of the enhancers showed only modest effect on their activity. These results are quite surprising since based on the analysis of endogenous genes in *zld-* mutant embryos in a previous study (Nien et al., 2011), we would have expected that the Zld binding site mutations would cause totally loss thee enhancer activities or cause large changes in the expression patterns of the reporter gene as described above. Based on our results, the effects observed in *zld-* mutant embryos may often be due to secondary effect. In support of this, has been shown previously for the otd_EHE enhancer that, while Zld binding site mutations did not seem to have effect on its activity, its expression pattern shifted posteriorly in *zld-* embryos (Xu et al., 2014).

Interestingly, even though the overall effects we observed as a result of Zld binding site mutations is modest for most of the enhancers, the observed effects often varied spatially and temporally. Previous studies suggested that one major function of Zld is to boost morphogen gradient activity (Foo et al., 2014; Hannon et al., 2017; Xu et al., 2014). Our results provide further support to this model. First of all, in consistence with the model, we showed that the Kr_D1, a class II Bcd activated enhancer active in the middle part of the embryo where Bcd concentration is low, is strongly dependent on Zld, similar to previous report (Xu et al., 2014). In addition, we showed that for both h_S1 and eve_S2, which belong to the class I BCD activated enhancers(Xu et al., 2014), in early c14 the expression in the posterior part of their expression stripe were more strongly affected by Zld binding site mutations, which made them the first examples in this class of enhancers to display spatially dependent Zld effect. Interestingly, three class I enhancers have been tested in *zld-* mutant embryo previously, but, counter intuitively, the only effect observed in all three cases was that their expression pattern shifted posteriorly, which probably due to indirect secondary effect resulting from loss of Zld in the mutant embryo (Xu et al., 2014). In addition to the Bcd dependent enhancers tested in this study, we also showed that the hb_stripe enhancer activated by Cad, also displayed spatially dependent effect, even though Cad does not form a gradient in older c14 embryo. This demonstrates that Zld can modulate the activity of target enhancers in different cells (or zone) in which the factor concentrations are different.

A comparison of h_1 and eve_S2 enhancers also provides interesting insight into Zld function. For eve_S2 enhancer, no effect from Zld binding site mutation was observed toward the anterior part of the embryo from ~30% egg length. However, for the h_S1 enhancer, which is normally expressed more anteriorly than eve_S2, with its stripe of expression centered around 27% egg length, significant effect from Zld binding site mutations was still observed around the center of the stripe. This suggests at a given position along the Bcd morphogen gradient, different enhancers can still display different Zld dependence; and some enhancers still use a combination of Zld sites and Bcd sites even if sufficient activation can be achieved by, for example, increasing the number of Bcd binding sites. It is possible in such cases, Zld may be required for other purposes.

One important phenomenon from our study is that the differences between the mutant and the wild type enhancers are generally greater at the beginning of c14 than in older embryos. While this is probably partly due to differences in the amount of transcripts accumulated in early mitotic cycles, a slower ramp up of the mutant enhancer activities at the start of c14 following mitosis from previous cycle provides another explanation. In support of the later explanation, we often see differences in the amount of nascent transcript signals in the nuclei of the wild type and mutant embryos (not shown). Such difference is particularly clear for the hb_stripe enhancer. We observed that, for this enhancer, the reporter signal was strong at the start of c14 for wild type enhancer and mainly located in the nuclei, while at this time no reporter signal was detectable for the Zld site mutant. A temporal delay in the expression of endogenous genes in *zld-* mutant embryo, or in the reporter activity driven by a Zld binding site mutated enhancer during early embryogenesis have previously been reported, even though an adequate explanation was lacking (ten Bosch, 2006; Nien et al., 2011). Our results suggest that the temporal delay may be due to the slow ramp up of enhancer activity following mitosis. Since before entering c14, the embryo is undergoing rapid mitotic cycle, a slow ramp up of transactivation following mitosis would prevent the target genes to be expressed given the short period of time the transcription process can occur during each mitotic cycle.

Taken together, our results show that Zld often exerts quantitative effect on enhancer activities, and that its effects on enhancers often vary temporally and spatially for a given enhancer. The early embryo development is a rapid process, with morphogens and spatially distributed activators, such as Cad, undergoing dynamic spatial and temporal changes. With its ability to boost activator activity, Zld can change the dynamics and spatial response of its target enhancers. Combined with the fact that different enhancers are associated different levels of Zld binding, Zld can exert a broad range of spatial-temporal effect on different enhancers, which may be important for generating the diverse expression patterns in the early gene network.

## Materials and Methods

### Plasmids and constructs

To create the reporter vector pLEP-BFYZ, the *eve* promoter in the pBΦY – lacZ reporter vector (Hare, 2008) was removed by restriction digestion using BglII, which cuts in the restriction enzyme polylinker region, and XhoI, which cuts at position +4 of the *eve* promoter sequence, and then the *eve* gene sequence from −860 to +4 was cloned in its place by Gibson assembly reaction. The *eve* fragment used in the Gibson assembly reaction was produced by the PCR amplifcation using the following primers (the upstream vector sequence is underlined): eve-860-F: 5’ GGCCAGATCCAGGTCGCAGCGGCCGCCTGGTCGAAGATCTCCTGGTACAGTTGGTACGCTGG 3’

Eve+40-R: 5’ TAACGAAGGCAGTTAGTTGTTGACTGTG 3’

The wild type enhancers tested were PCR amplified from genomic DNA, while the Zld site mutants were synthesized by GenScript (GenScript.com). The sequences of the wild type and mutant enhancer sequences, including information of the chromasome coordinates of the enhancers, are provided in the supplement. Both the wild and mutant enhancers were cloned into the NotI and BglII sites of the pLEP-BFYZ.

The Zld binding site mutations in the *eve* stripe 2 enhancer region in the eve BAC was created through recombineering (Lee et al., 2001; Liu et al., 2003; Warming et al., 2005; Venken et al., 2009). The starting eve BAC plasmid P[acman]-CH322-103k22-GFP, which contains the eGFP-Kan^r^ tag sequence at the C-terminus of the *eve* CDS, was previously described (Venken et al., 2009). We removed the CDS of the *eve* gene in this construct to generate peve-GFP by standard recomibineering procedure, by first replacing the *eve* CDS with galK sequence amplified from a galK plasmid pBALU1 using primers flanked by *eve* sequences:

eve-CDS-galK-F: 5’

<u>TTAATATCCTCTGAATAAGCCAACTTTGAATCACAAGACGCATACCAAAC</u>CCT GTTGACAATTAATCATCGGCA 3’

eve-CDS-galK-R: 5’

<u>GCGCCCTGAAAATAAAGATTCTCGCTTGCAGTAGTTGGAATATCATAATC</u>TCA GCACTGTCCTGCTCCTT 3’.

Then the galk sequence was replaced with:

5’ ATATCCTCTGAATAAGCCAACTTTGAATCACAAGACGCATACCAAACATGGATTATGATATTCCAACTACTGCAAGCGAGAATCTTTATTTTCAGGGCGC 3’

To create the *eve* stripe 2 Zld site mutations, the *eve* stripe 2 sequence was first replaced with galK sequence through recombineering using PCR products generated with the following primers:

eve-s2-galK-F: 5’ AGGGATTCGCCTTCCCATATTCCGGGTATTGCCGGCCCGGGAAAAT

GCGACCTGTTGACAATTAATCATCGGCA 3’

eves2-15-galKr: 5’ TTGCGCAAGTTAGCTTGGAGGTTTGGCCAAAAAAATCGTGGGGTCC

ACCCTCAGCACTGTCCTGCTCCTT 3’

Then, the galK sequence was replaced with the mutant stripe enhancer containing the Zld site mutations, which was synthesized as genBlock (Integrated DNA technologies).

### Transgenic Flies

The transgenes were generated by injecting the reporter construct plasmids into the embryos from flies that carry the VK33 (Venken et al., 2006) attp landing site on chromasome 3 using the services of Bestgene Inc. Homozygous lines for the transgenes were established by selfing the balanced red eye flies obtained from Bestgene Inc following the injection.

### Florescence *in situ* hybridization and embryo imaging

FISH was carried out with embryos collected from the transgene flies and fixed with formaldehyde as previously described (Kosman et al., 2004) with some modifications.

To prepare the RNA probe for the reporter gene, the eGFP reporter gene and the kan^R^ gene sequences were amplified by PCR separately from the *eve* reporter construct using the following oligos:

eGFP-F: 5’ GGATTATGATATTCCaACTACTGCAAGCGAG 3’

T7p-eGFP-R: 5’ CAGGTCTGAGTAATACGACTCACTATAGGGTACAGCTCGTCCATGCC

GAG 3’

Kan-F: 5’ GTTACTGGCCGAAGCCGCTTG 3’

T7p-Kan-R: 5’ CAGGTCTGAGTAATACGACTCACTATAGGGAAGAACTCGTCAAGAAGGC

GATAGAAG 3’

The reverse primers used contained T7 promoter sequence at their 5’ end, allowing the PCR products to be used directly for RNA probe synthesis. The probes were made using Thermo Transcript Aid T7 High Yield Transcription Kit (ThermoFisher Scientific, cat. no. K0441) with the Digoxigenin(DIG) RNA labeling mix from Roche (cat. no. 1277073) and were not hydrolyzed before being used.

The embryos were collected for 2 hours, aged for 1 hr and 20 min and were fixed as described (Kosman et al. 2004). The preparation before hybridization and the hybridization step were carried out as described except that the protease K treatment and the second post-fix step that followed were skipped. For florescent detection, the embryos were incubated with pre-absorbed sheep anti-DIG-POD (Roche, cat. no. 11207733910). Following incubation and washing, the embryos were mixed and incubated with TSA coumarin in amplification buffer and incubated by rocking. The embryos were washed, and near the end of wash were incubated with Sytox Green nucleic acid stain (ThermoFisher Scienctific, cat. no. S7020) to label the nuclei. The embryos were washed again once, and mounted in 70% glycerol, 2.5% DABCO.

The embryo imaging was performed on a Zeiss LSM800 microscope with a 10x objective. For all experiments, Z stack images were taken from top to the mid-section of each embryo with the image size set at 2014 × 849, or a resolution of about 0.27 µm/pixel. The other settings differed between experiments, which was not anticipated to affect the results since reporter activities detected for the mutant enhancers were always compared to the signals detected for wild type enhancer in the same experiment. For earlier experiments, 15 z stacks were taken for each embryo with thickness of about 5.6 µm, and the fluorescence signals for coumarin with an excitation wavelength of 405 and Sytox Green with excitation wavelength at 488 were obtained sequentially with the “smart” setting. For later experiments, 30 z stacks of images per embryo were taken, and the two fluorescence signals were obtained simultaneously with the “fast” setting. In all cases, the laser power and digital gain for each dye were adjusted in each experiment to make sure the maximum signal obtained was below saturation.

The images were analyzed with the Zeiss image analysis software zen 2. A 2-D image was generated for each embryo from the z stack images using the orthogonal maximum projection function. Then the fluorescent signal profiles for the RNA (coumarin) and nuclei (sytox green) were generated for a rectangle area drawn to the length of the embryo along the A/P axis, and with a width of 200 pixels, about a third of the embryo width. The signals along the A/P axis were then divided into 100 bins corresponding to the percent egg length. The coumarin signals were then divided by the corresponding Sytox signals as a simple way of normalization to correct for the nuclear density variation between different part of the embryo and between embryos. Furthermore, since the reporter signal between egg length 50% - 60% is the lowest in the coumarin signal profile, and not affected by the presence hb DAE, and the average of signal in this region was taken as background and subtracted from the signal throughout the whole embryo.

We carefully staged the embryos so that the dynamics in the changes of the reporter activity in the mutant and wild type embryos can be compared. We divided the embryos into four stages, c14-(A–D), based on the extent of cell membrane invagination, and further divided c14-A into c14-A0, A1 and A2 based on nuclear shape: c14-A0 is when the nuclei have mostly just finished cycle 13 mitotic cycle and have become round, c14-A1 is when the nuclei have expanded but not elongated, while c14-A2 is after the nuclei have elongated and the cell membrane invagination is about a quarter complete. For each experiment, the signal profiles of different embryos in each age group were combined to generate the average profiles, which are shown in the figures.

---

**Fig. S1.**
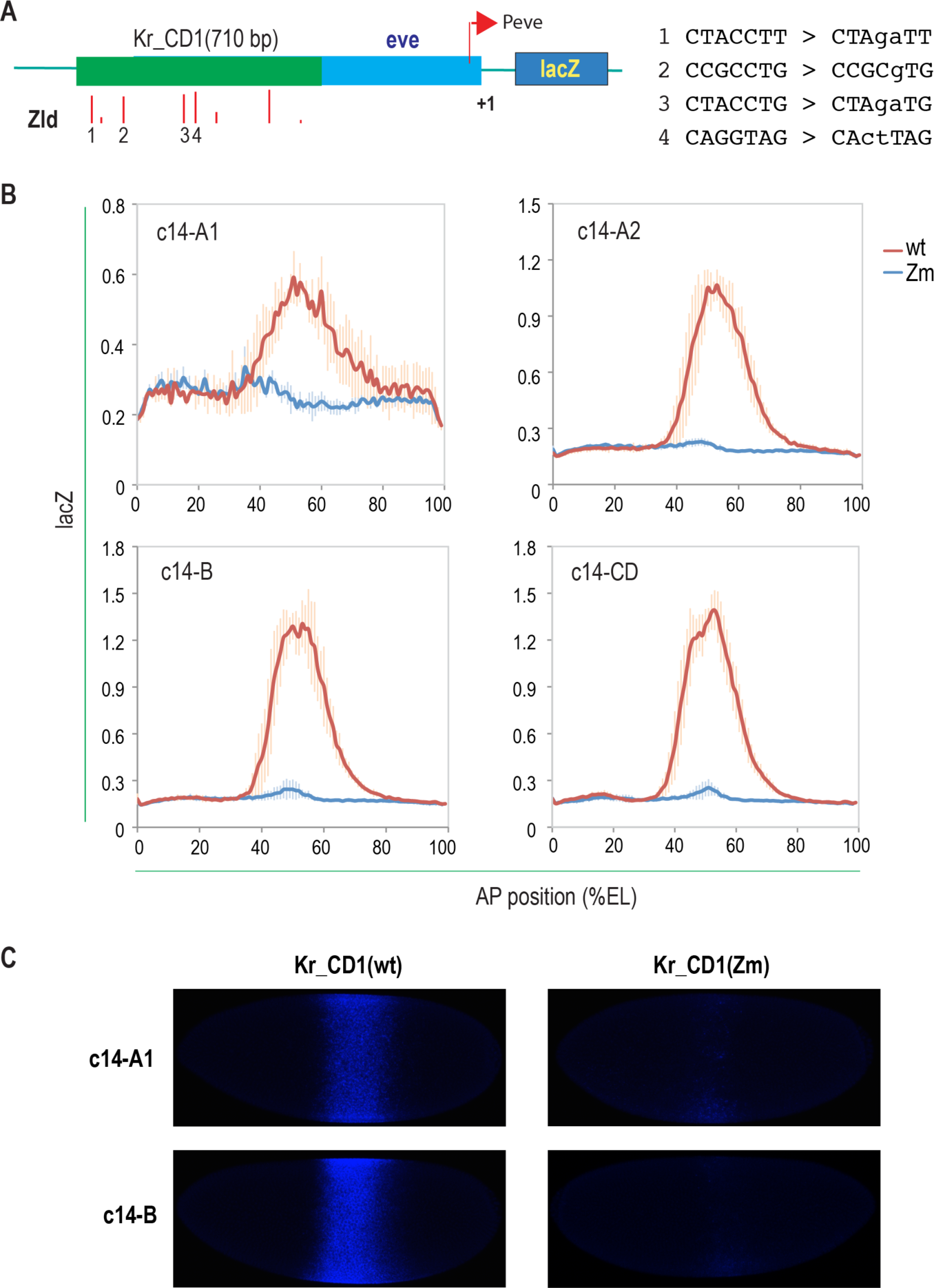
Effect of Zld binding site mutations on Kr_CD1 enhancer at different stages. The same as Fig. 1B-D, except results from more stages are shown. A. Schematic of the Kr_CD1 enhancer. Zld site mutations are also shown. B. the reporter activities, detected and analyzed as described in Fig. 1B, at different stages of the embryos are shown. C. representative images of wild type and mutant embryos

**Fig. S2.**
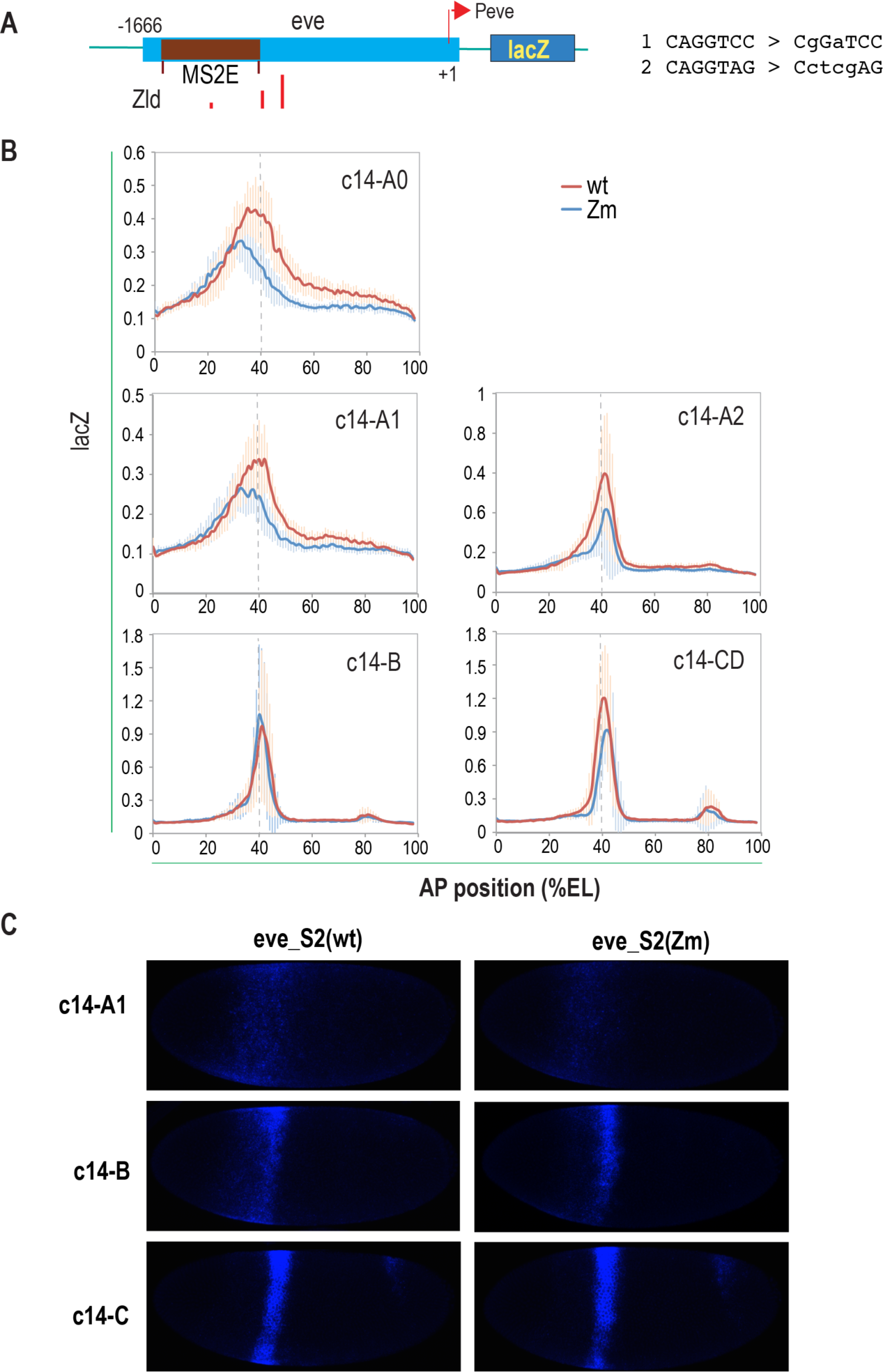
Effect of Zld binding site mutation on eve_S2 enhancer at different stages. The same as Fig. 1E-G, except results from more stages are shown. A. Schematic of the eve_S2 enhancer construct, which contains the *eve* sequence from −860 to −1666 overlapping previously defined minimal stripe 2 enhancer (MS2E), in addition to the −860 - + 163 *eve* sequence present in the standard pLEP-PFYZ vector. Zld site mutations are also shown. B. the reporter activities, detected and analyzed as described in Fig. 1B, at different stages of the embryos are shown. C. representative images of wild type and mutant embryos

**Fig. S3.**
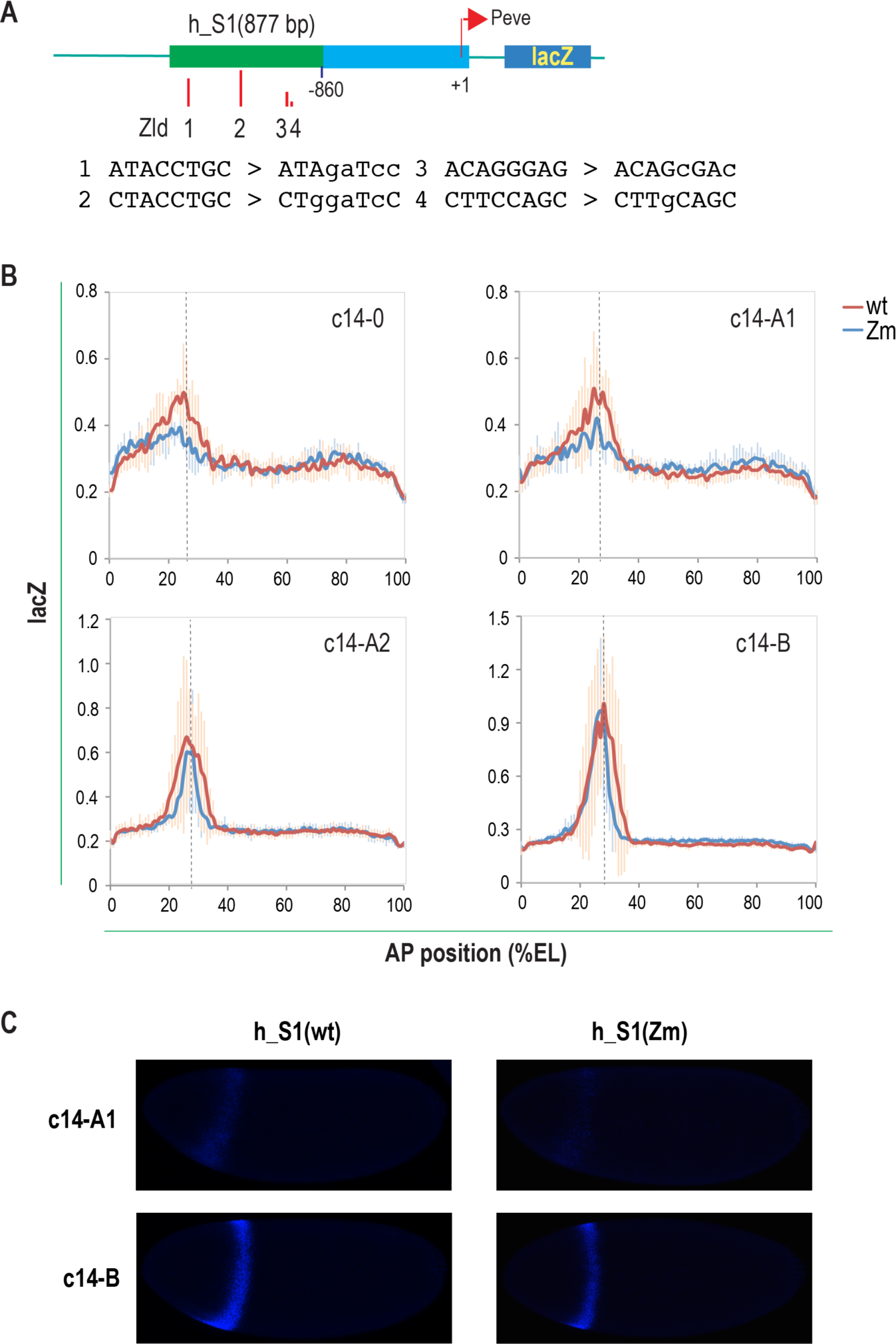
Effect of Zld binding site mutations on the h_S1 enhancer at different stages. The same as Fig. 1H-J, except results from more stages are shown. A. Schematic of the h_S1 enhancer. Zld site mutations are also shown. B. the reporter activities, detected and analyzed as described in Fig. 1B, at different stages of the embryos are shown. C. representative images of wild type and mutant embryos

**Fig. S4.**
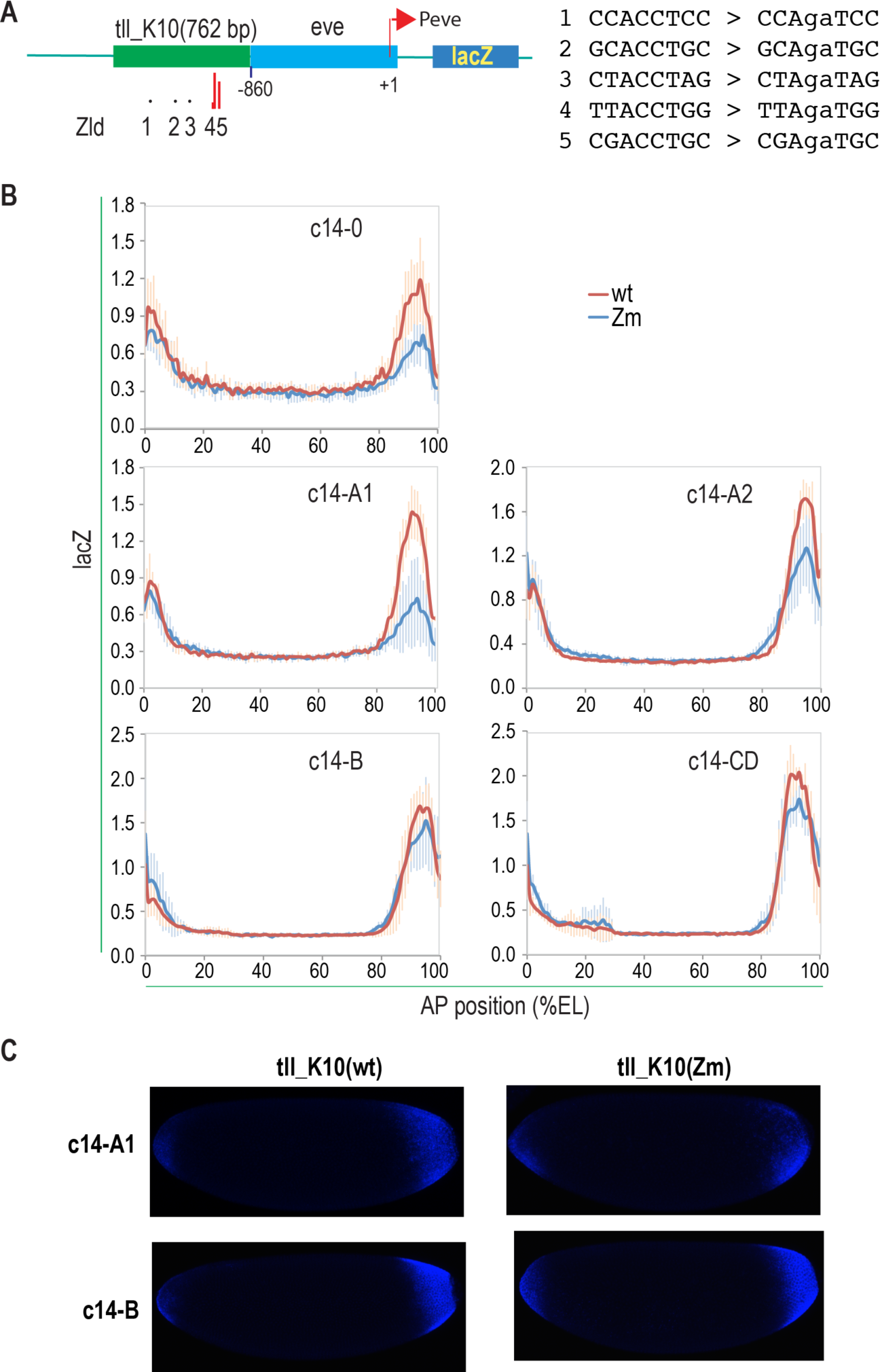
Effect of Zld binding site mutation on tll_K10 enhancer at different stages. The same as Fig. 2A-C, except results from more stages are shown. A. Schematic of the tll_K10 enhancer. Zld site mutations are also shown. B. the reporter activities, detected and analyzed as described in Fig. 1B, the embryos of different stages are shown. C. representative images of wild type and mutant embryos

**Fig. S5.**
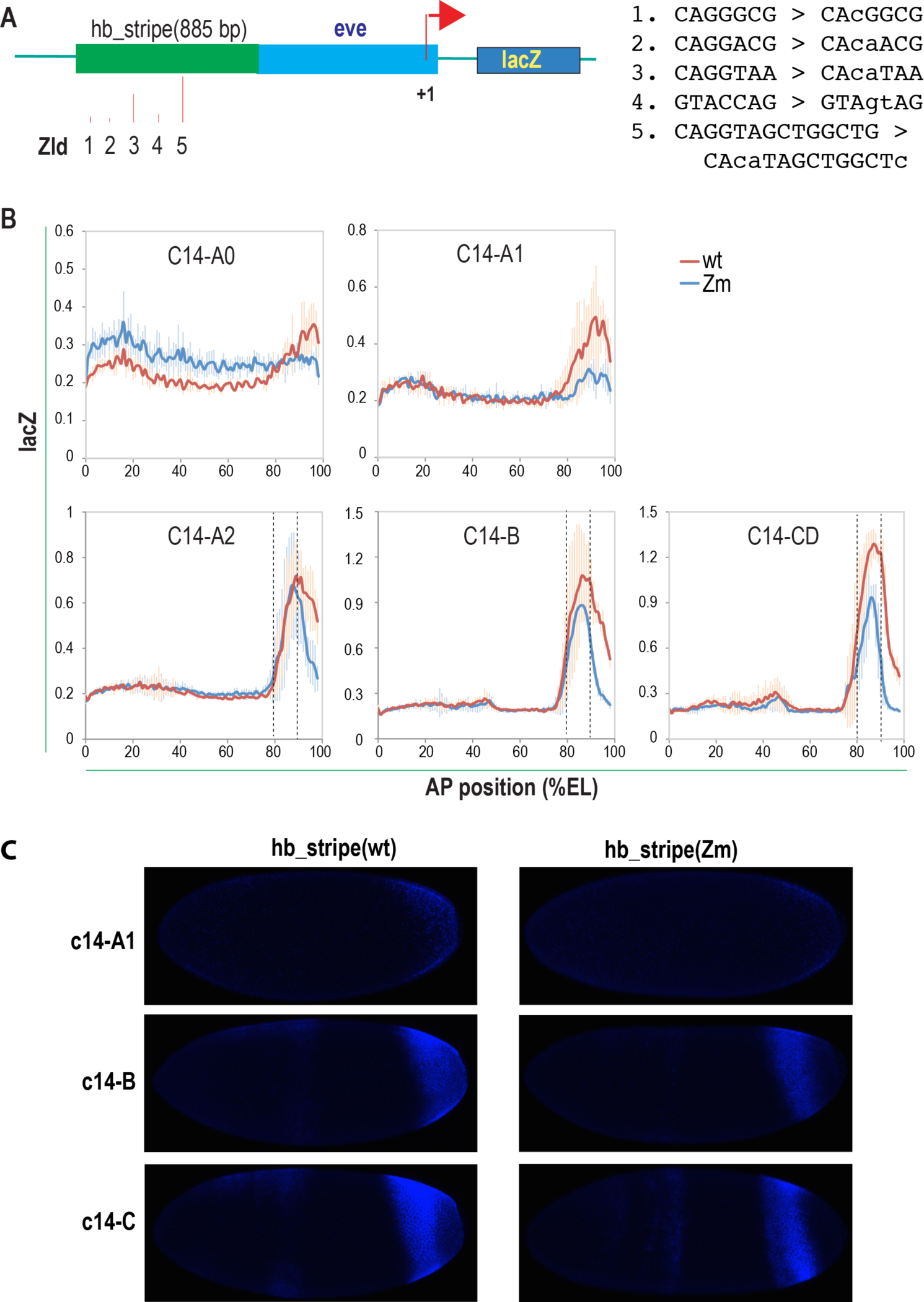
Effect of Zld binding site mutations on the hb_stripe enhancer activity at different stages. The same as Fig. 2D-F, except results from more stages are shown. A. Schematic of the hb_stripe enhancer. Zld site mutations are also shown. B. the reporter activities, detected and analyzed as described in Fig. 1B, at different stages of the embryos are shown. C. representative images of wild type and mutant embryos

